# Growth and eGFP-production of CHO-K1 suspension cells cultivated from single-cell to lab-scale

**DOI:** 10.1101/2021.05.20.444654

**Authors:** Julian Schmitz, Oliver Hertel, Boris Yermakov, Thomas Noll, Alexander Grünberger

## Abstract

Scaling down bioproduction processes became a major driving force for more accelerated and efficient process development over the last decades. Especially expensive and time-consuming processes like the production of biopharmaceuticals with mammalian cell lines benefit clearly from miniaturisation, due to higher parallelisation and increased insights while at the same time decreasing experimental time and costs. Lately, novel microfluidic methods have been developed, especially microfluidic single-cell cultivation (MSCC) devices proofed to be valuable to miniaturise the cultivation of mammalian cells. So far growth characteristics of microfluidic cultivated cell lines were not systematically compared to larger cultivation scales, however validation of a miniaturisation tool against initial cultivation scales is mandatory to proof its applicability for bioprocess development. Here, we systematically investigate growth, morphology, and eGFP-production of CHO-K1 cells in different cultivation scales including microfluidic chip (230 nL), shake flask (60 mL), and lab-scale bioreactor (1.5 L). Our study shows a high comparability regarding growth rates, cellular diameters, and eGFP production which proofs the feasibility of MSCC as miniaturised cultivation tool for mammalian cell culture. In addition, we demonstrate that MSCC allows insights into cellular heterogeneity and single-cell dynamics concerning growth and production behaviour which, when occurring in bioproduction processes, might severely affect process robustness. Eventually, by providing insights into cellular heterogeneity, MSCC has the potential to be applied as a novel and powerful tool in the context of cell line development and bioprocesses implementation.

## Introduction

The number of biotechnologically manufactured products like biopharmaceuticals increased rapidly over the last decades (Walsh 2018). Consequently, there is a continuing desire for new and more efficient bioprocesses to cover the increasing demand. Lately, the development of improved bioprocesses went hand in hand with the technological progress of miniaturisation (Hemmerich et al. 2018). Since first approaches the focus of scale-down applications lies on the same ambitions: Minimise costs, reduce experimental time, and simultaneously increase insights (Janakiraman et al. 2015; Kim et al. 2012). Given that mammalian cell culture processes require considerably longer experimental time spans than bacterial processes, and process development is often based on empirical testing of multiple interdependent parameters (Neubauer et al. 2013), especially time reduction and increasing experimental throughput are highly desirable to enhance process development (Rameez et al. 2014; Zhang et al. 2010). Furthermore, maximising the analytical throughput and expanding the degree of parallelisation improves not only process development but also cell line or medium design (Betts and Baganz 2006).

Miniaturising a bioproduction process often depends on novel bioreactor concepts that do not match the original cultivation conditions or cultivation vessel geometry of the manufacturing scale which makes systematic validation mandatory. Therefore, to qualify a technology for miniaturisation, the recorded data needs to be verified against data from original scale approaches (Li et al. 2006; Tsang et al. 2014). If the generated data are not comparable, a prediction for the original scale based on the data from the miniaturised scale is unfeasible and ultimately leads to deviations in process development or challenges in eventual scale-up (Bertrand et al. 2018; Betts and Baganz 2006).

Lately, different approaches to miniaturise mammalian cultivation for bioprocesses development and screening have been introduced, all based upon different concepts ranging from shake flask applications to shaken microtiter plates and miniaturised stirred bioreactors (Zhang et al. 2010). An already established miniaturised stirred bioreactor consist in the ambr™ platform, which proved suitable to emulate temperature, dissolved oxygen and pH profiles matching large scale bioreactors and shows comparable growth and productivity (Rameez et al. 2014). Using a shaken 24-well single-use cassette with bubble columns, Betts et al. were able to mimic industrially relevant fed-batch processes with comparable growth and production performance (Betts et al. 2014). Furthermore, orbitally shaken tubes have been applied as miniaturisation tool to optimise operating conditions of mammalian perfusion cultures and showed good comparability with benchtop bioreactors (Wolf et al. 2018). Besides already established approaches, microfluidic cultivation tools became increasingly relevant in terms of downsizing and mark the next level of miniaturisation (Marques and Szita 2017; Mehling and Tay 2014). On the one hand microfluidic approaches can be applied to investigate one discrete bioprocess related question, on the other hand microfluidics can be applied to miniaturise the whole bioprocess (Bjork et al. 2015).

In addition to the already mentioned preferences of miniaturisation, microfluidics additionally extends the toolbox of already established approaches by the feature to cultivate and analyse cells with single-cell resolution (Hung et al. 2005; Kolnik et al. 2012; Lindström and Andersson-Svahn 2010). Therefore, microfluidic single-cell tools can be applied to analyse cellular heterogeneity which would stay masked by standard average measurements conventionally used in bioprocess research. Due to the genetic plasticity and origin of every industrial production cell line, utilised populations doubtlessly exhibit genetic or phenotypic heterogeneity (Barnes et al. 2003; Pilbrough et al. 2009) which has been ignored in bioprocess development over decades (Schmitz et al. 2019).

In the context of bioprocess research and development, microfluidic single-cell cultivation (MSCC) represents the tool of choice to investigate cellular behaviour concerning heterogeneity in growth and morphology, proliferation, and productivity (Schmitz et al. 2019). In contrast to other single-cell analysis applications like flow cytometry or droplet microfluidics, MSCC combines features necessary for long-term cultivation under controlled environmental conditions, that are needed for bioprocess near research with analytical prospects like live cell imaging and thereby facilitate high spatio-temporal resolution of cellular behaviour (Grünberger et al. 2014).

In this work we present a comparative study with a focus on growth and production of CHO-K1 cells at different scales, ranging from microfluidic single-cell cultivation (230 nL) and shake flasks (60 mL) to benchtop bioreactors (1.5 L). Lately we introduced a platform for mammalian single-cell cultivation which, for the first time, enabled cultivation of mammalian suspension cell lines with single-cell resolution under process near environmental conditions (Schmitz et al. 2020). To approach the question if mammalian single-cell cultivation is feasible for miniaturising cultivation for future bioprocess screening approaches, we here investigate possible differences in terms of growth behaviour, cell morphology, and eGFP-production. Furthermore, we give an example of how MSCC can enlarge the insights into single-cell dynamics.

## Material and Methods

### Cell culture and medium

In this work the CHO-K1 cell line ATCC CCL-61, adapted to growth in suspension, was applied as model for other mammalian cell lines used in biotechnology. Furthermore, an eGFP-producing CHO-K1 pool was applied for comparative studies of production behaviour between the cultivation scales. Therefore, the same CHO-K1 cell line was transfected with a vector containing eGFP under the control of the endogenous HSPA5 promoter and a puromycin resistance for selection (Fig. S1). Two weeks after transfection and cultivation with 8 µg/mL puromycin, the eGFP gene was randomly integrated into the genome and the heterogenous cell pool was cryopreserved.

For cultivation, the commercially available medium (TCX6D, Xell, Germany), supplemented with 6 mM glutamine was utilised. The pre-culture of eGFP expressing CHO-K1 cells was further supplemented with 8 µg/mL puromycin but the main cultivation was executed without selective pressure. Initial CHO cell culture was inoculated from a uniform working cell bank and cultivated at a temperature of 37 °C, 5 % CO_2_, 80 % humidity, and 185 rpm (maximal deflection 50 mm) on an orbital shaker (ES-X, Kühner AG, Switzerland). For reproducibility, the pre-culture was passaged in the exponential phase between two and three times before starting any of the following cultivation experiments.

### Microfluidic single-cell cultivation

Microfluidic single-cell cultivation was performed as described previously (Schmitz et al. 2020). The employed PDMS-glass-device was mounted onto an automated inverted microscope for phase contrast microscopy (Nikon Eclipse Ti2, Nikon, Germany). For seeding cells into the cultivation chambers, CHO-K1 cell suspension was manually flushed through the cultivation device until the cultivation chambers were filled with cells sufficiently. Next, fresh medium mixed with conditioned medium, obtained from exponential growth phase, in a ratio of 1:1 to simulate substrate and metabolite situation in standard batch cultivations was constantly perfused through the supply channels by low pressure syringe pumps (neMESYS, CETONI, Germany) with a flow rate of 2 µL/min. Constant cultivation conditions of 37 °C and 5 % CO_2_ were controlled by microscope an incubator systems (Cage incubator and H201-K-FRAME GS35-M, OKO Touch, Okolab S.R.L., Italy). For microscopic analysis of single cells, a 40x objective was applied and relevant positions were analysed every 20 min (NIS Elements AR 5.20.01 Software, Nikon Instruments, Germany).

### Shake flask cultivation

Shake flask cultivation was performed as triplicates in 125 mL shake flasks (Flat Base, TriForest, USA) with a cultivation volume of 60 mL, each inoculated at 5×10^5^ cells/mL from one 250 mL pre-culture to assure reproducibility. Cultivation temperature, CO_2_ atmosphere, and humidity were matching the pre-culture. Every 12 hours samples for analysis of growth, viability, and morphology were taken and measured using CEDEX cell counter (Innovatis, Germany).

### Bioreactor cultivation

Bioreactor cultivation was performed as duplicates in 2 L Biostat B-DCU bioreactors (Sartorius AG, Germany) with a working volume of 1.5 L, inoculated at 5×10^5^ cells/mL. The cultivation temperature, pH-value and dissolved oxygen concentrations were controlled at 37 °C, at 7.2, and 40 % of the air saturation. Stirring speed was set to 150 rpm using a Rushton turbine.

### Growth rate analysis

As key indicator for growth comparison, the growth rate µ_max_ of each cultivation was analysed. For microfluidic single-cell cultivation, the cell number was determined every 12 h by analysing time-lapse images. By offsetting the cultivation chamber’s volume of 3.2 × 10^−7^ mL against the number of cells inside, the enumerated cell number can be converted into a cell density. To evaluate growth rates, cell densities from microfluidic cultivation and viable cell densities from shake flask and bioreactor cultivation were plotted against cultivation time semi-logarithmically to identify the relevant interval for µ_max_ determination. In the following, µ_max_ was determined graphically from the slope of the exponential growth phase of each plot using OriginPro (OriginPro 2020 9.7.0.188, OriginLab Corporation, USA).

### Cell morphology analysis

In order to compare cell morphology over scales, for microfluidic cultivation cellular area was determined manually using ImageJ 1.52p (Schindelin et al. 2012) and converted into cell volume by multiplying cell areas with the cultivation chamber height of 8 µm, implying a cylindric cellular shape inside the microfluidic device. Assuming a natural sphere-shaped cellular morphology without the restrictive height, calculated volumes were again converted into cellular diameter data. To enlarge sample size and thereby statistical significance, the number of analysed cultivation chambers was increased by six randomly selected cultivation chambers to a total number of nine cultivation chambers. For shake flask and bioreactor, cellular diameters were determined via CEDEX simultaneously with cell density and viability measurements.

### Fluorescence analysis

eGFP fluorescence was analysed by flow cytometry measurements for bioreactor and shake flask cultivation applying S3e™ Cell Sorter (Bio-Rad, Germany) applying a 488 nm laser in combination with a 525/30 nm filter. The obtained data was analysed and visualised using FlowJo (Becton, Dickinson and Company, USA). For microfluidic cultivation fluorescence microscopy was utilised with an exposure time of 20 ms and 10 % intensity. Subsequently, 16-bit TIFF images for relevant time points were created to ensure maximal information density by the amount of grey scale values and a cell’s mean grey scale value was determined manually as arbitrary unit using ImageJ 1.52p (Schindelin et al. 2012) to describe cellular fluorescence level.

## Results and Discussion

### Key characteristics of bioreactor, shake flask, and MSCC

In the presented study the growth characteristics of CHO-K1 cells in three different cultivation setups namely bioreactor, shake flask, and microfluidic single-cell cultivation device (Fig. 1a) were compared. Besides the variance in cultivation volume ranging from nanolitre to litre, the cultivation setups used within this study differ in their material and their mixing approach (Fig. 1b). While the here utilised bioreactors consist of glass vessels, shake flasks are made of polycarbonate and the applied microfluidic device is a hybrid of glass and PDMS. Therefore, cells experience different surface interactions which might influence their physiology and thus growth behaviour. Likewise, mixing is achieved by different mechanisms namely stirring (bioreactor), shaking (flask), and diffusion (MSCC) which might have an influence on possibly arising environmental gradients or temporary limitations in oxygen saturation or nutrient concentration.

**Fig. 1:**
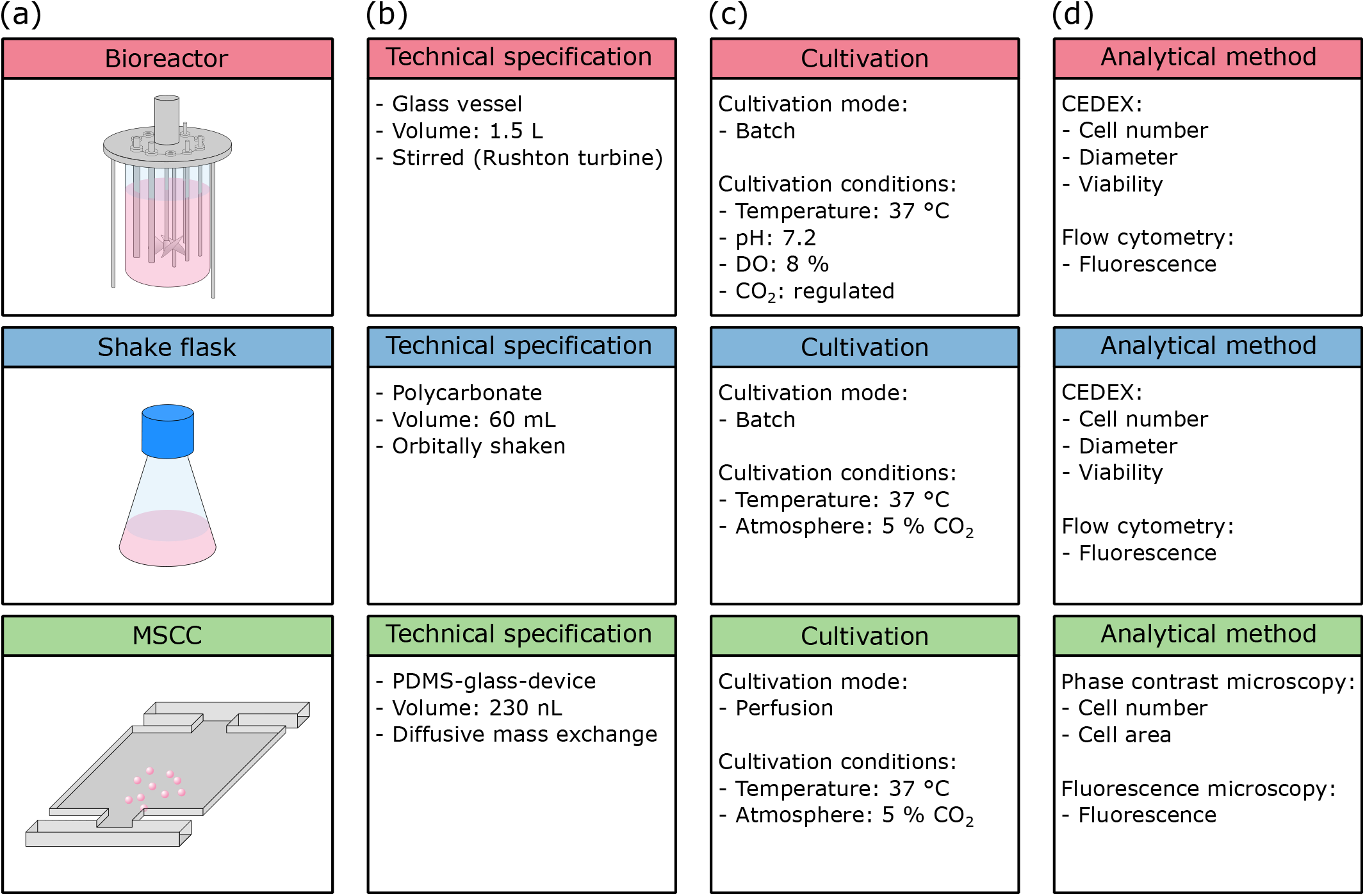
Overview of the cultivation setups used within this study. (a) Schematic figure of the cultivation setups, (b) technical specifications, (c) cultivation conditions and mode, and (d) analytical methods.

Bioreactor and shake flask cultivations were executed in batch mode whereas MSCC is performed as perfusion and thus ensuring constant environmental conditions over the whole cultivation course (Fig. 1c). Due to integrated process analytical technology, cultivation conditions inside the bioreactor are feedback regulated while conditions inside the shake flask and microfluidic device are not subjected to any active control loops besides a constant cultivation temperature and CO_2_ atmosphere.

On analytical level (Fig. 1d), bioreactor and shake flask again share the same procedures: Cell number for growth analysis and diameter examination to address cellular morphology are performed off-line by applying a CEDEX cell counter. Additionally, the viability of the analysed sample can be detected. To investigate production behaviour, here in form of eGFP fluorescence, flow cytometry measurements are conducted. In contrast, for microfluidic cultivation cell number and cellular area are determined from phase contrast microscopy as well as fluorescence level from fluorescence microscopy. Viability cannot be determined precisely in MSCC since distinguishing between viable and dead cells is only possible if cell death has a noticeable influence on morphology which only occurs unreliably for the here cultivated CHO-K1 cells.

### Comparison of growth behaviour

Establishing a new miniaturisation approach like MSCC requires systematic testing of growth behaviour between traditional cultivation scales and miniaturised scales to proof its comparability. Therefore, we analysed the growth of CHO-K1 cells with a focus on growth progression and µ_max_. The following experiments were all realised with identical cultivation medium, started from the same master cell bank, and were inoculated after a uniform pre-culture proceeding to guarantee comparability between different scales.

The curve progressions of the (viable) cell densities illustrated in Figure 2 are very similar between bioreactor, shake flask, and microfluidic device. Initial exponential growth from inoculation until three days of cultivation resembles each other in appearance. Since shake flask and bioreactor cultivations were performed in batch-mode, following on the exponential phase growth rate declines (Fig. S2). In contrast, growth behaviour of cells cultivated in the microfluidic device stays constantly exponential until the cultivation chamber is entirely filled (Fig. S2, Video S1), due to optimal nutrient supply owing to continuous medium perfusion. Considering the limited cultivation chamber volume, microcolonies inside the microfluidic device reach about 30-times higher maximal cell densities with approx. 4.5×10^8^ cells/mL compared to bioreactor and shake flask cultivation. Supplemental analysis of cell viability during bioreactor and shake flask cultivation reveals no significant deviation from expected trends (Fig. S3).

**Fig. 2:**
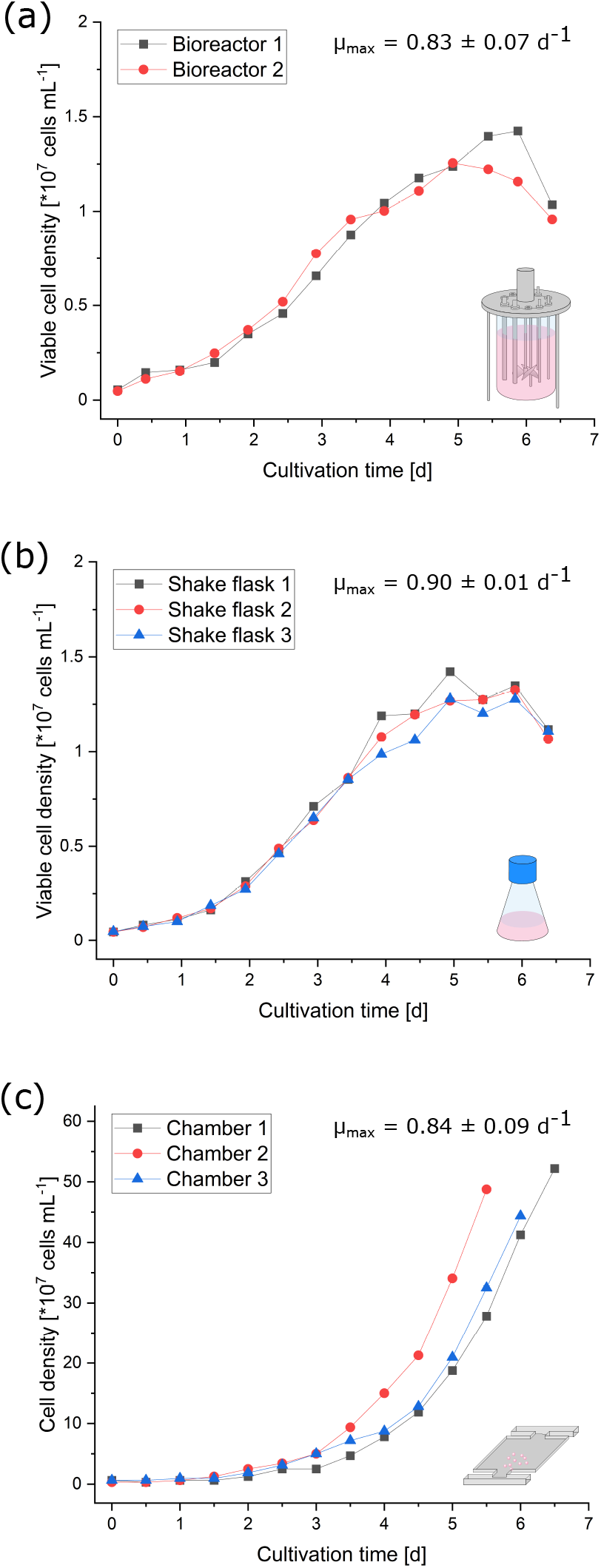
Growth comparison of the different cultivation scales. (a) Viable cell density of two parallel bioreactors plotted against the cultivation time. (b) Viable cell density of three parallel shake flasks plotted against the cultivation time. (c) Cell density of three cultivation chambers plotted against the cultivation time.

Comparing respective maximal growth rates ranging from 0.83 ± 0.07 d^-1^ in bioreactors, over 0.84 ± 0.09 d^-1^ with MSCC, to 0.90 ± 0.01 d^-1^ in shake flasks also indicates a reliable comparability of miniaturised CHO-K1 cultivation and classic cultivation in millilitre and litre scale.

### Comparison of cellular morphology

Besides growth behaviour, also cellular morphology during MSCC needs to be validated regarding to shake flask and bioreactor. As a reference value single-cell diameter and its distribution within the analysed sample was investigated. Considering possible morphological changes over the course of cultivation, cell diameter distribution was measured at three time points during cultivation considering the respective growth phases in bioreactor and shake flask cultivation (Fig. 2): Straight after inoculation (t = 0 d), after three days (t = 3 d) at the end of exponential growth phase, and after five days (t = 5 d) at the beginning of stationary phase.

As can be seen in Figure 3, the cellular diameters of the analysed samples from MSCC, shake flask, and bioreactor show nearly identical distributions with a main peak around 14 µm. However due to the relatively small sample size for microfluidical data at t = 0 d, resulting from seeding only one to three cells into every cultivation camber, the statistical significance of the illustrated distributions from shake flask and bioreactor is more reliable (Fig. 3 left).

**Fig. 3:**
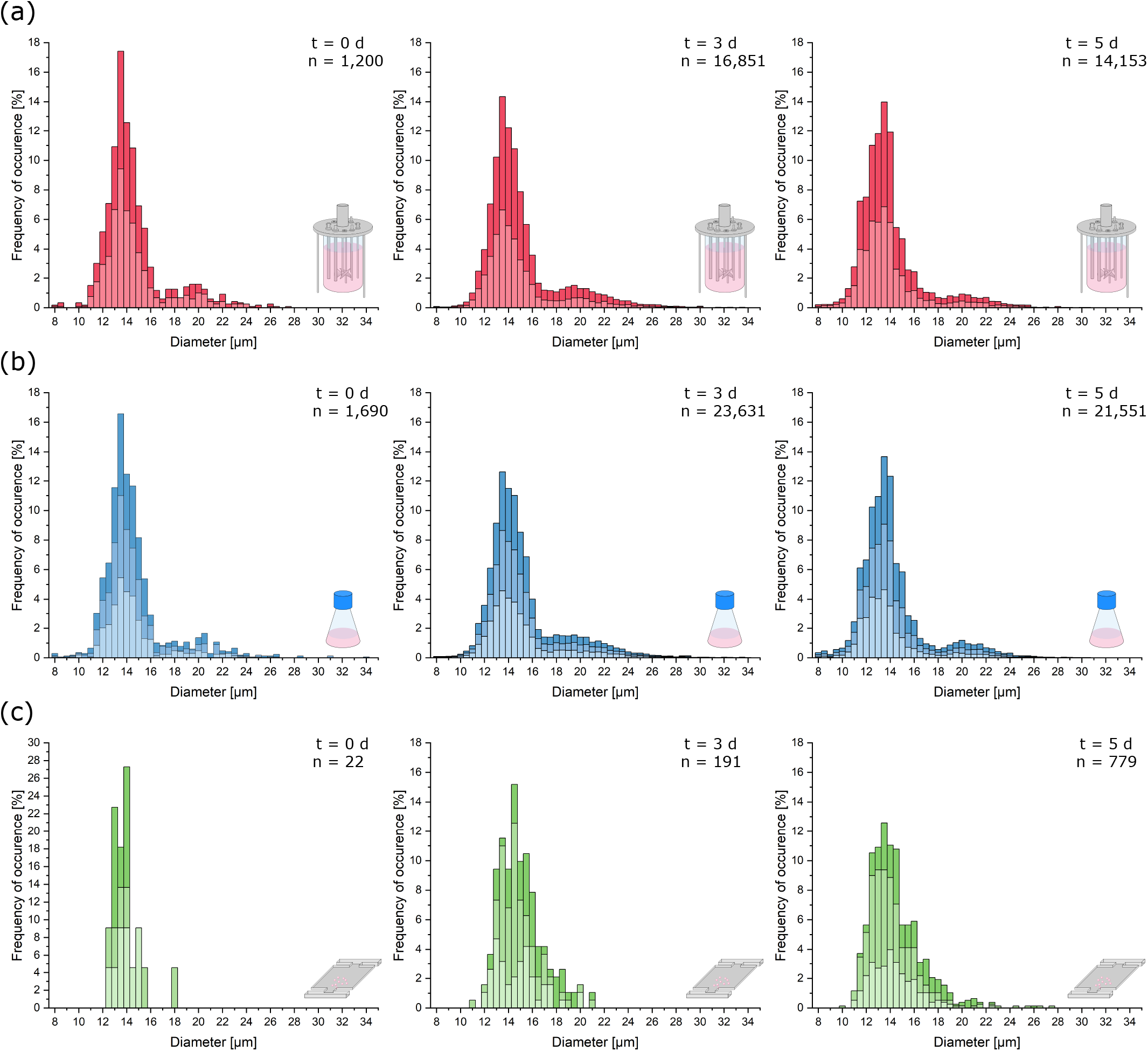
Cell morphology comparison of the different cultivation scales. Relevant cellular diameters are plotted against the frequency of their occurrence cumulated from the respective replicates of the analysed samples. (a) Cellular diameter distribution of the bioreactor cultivation right after inoculation (t = 0 d), after 3 days of cultivation (t = 3 d), and after 5 days (t = 5 d). (b) Cellular diameter distribution of the shake flask cultivation right after inoculation (t = 0 d), after 3 days of cultivation (t = 3 d), and after 5 days (t = 5 d). (c) Cellular diameter distribution of the microfluidic cultivation right after inoculation (t = 0 d), after 3 days of cultivation (t = 3 d), and after 5 days (t = 5 d).

After three days of cultivation the distributions of single-cell diameters from bioreactor and shake flask show more cells with a diameter above 18 µm compared to the microfluidic cultivation (Fig. 3 middle). For shake flask and bioreactor, the distribution of bigger cells becomes more uniform compared to the inoculum.

After 5 days the distribution of the microfluidic cultivation narrows further around a mean cellular diameter of 14 µm with a plateau at diameters between 14.5 and 16 µm. In shake flask more cells with a diameter below 11 µm can be observed and the overall distribution slightly hints at a second less dominant peak with cell diameters between 19 and 22 µm (Fig. 3b right). The same process can be hypothesised for the microfluidic cultivation (Fig. 3c right). The diameter data of the bioreactor shows the same characteristics as the data obtained from shake flask cultivation (Fig. 3a right). In comparison to the distribution from exponential growth phase, the portion of cells with a diameter over 18 µm decreases.

In general, cellular diameter data obtained from MSCC shows the same distribution and trends over the course of cultivation as diameters determined from shake flask and bioreactor cultivations with only minor differences. Thus, the limited cultivation chamber height seems not to influence the morphological characteristics of the analysed cells. The analysis of the cells’ growth behaviour already showed that growth rate is not affected by the restricted device dimensions. Morphologically, a potential shift and thereby adaptive behaviour over the cultivation’s course from normal diameters around 12 to 14 µm towards diameters of 8 µm equalling the cultivation chamber height is not noticeable.

### Green fluorescent protein production

The most important parameter within bioproduction represents the cells’ productivity. As model product eGFP was chosen to analyse cellular productivity. Therefore, an eGFP synthesising CHO-K1 cell pool was cultivated following the same protocols already established for the previous experiments.

Conventionally, fluorescence behaviour of cells cultivated in bioreactor and shake flask is analysed applying flow cytometry. For this reason, we determined the fluorescence signal of bioreactor and shake flask samples, taken every 24 h, with help of S3e™ Cell Sorter. Figure 4 shows the distribution of fluorescence intensity of the analysed cells ranging from inoculation to death phase after six days of cultivation (see Fig. S4 for growth curves). As can be seen, both bioreactor and shake flask show a broad distribution of fluorescent cells. For illustrative reasons non-fluorescent cells, which are present at any time during the cultivation, are not displayed here but can be seen in Figure S5. Corresponding values are listed in Table S1. For the data obtained from the bioreactor a clear tendency towards higher fluorescence intensities can be seen (Fig. 4a, Tab. S1). In general, the whole distribution gets broader and shifts towards higher fluorescence intensities. During shake flask cultivation the fluorescence distribution of the analysed cells does not change its character as observed for the bioreactor cultivation (Fig. 4b) and the mean fluorescence intensity stays unchanged (Tab. S1). For both cultivation methods the ratio of fluorescent to non-fluorescent cells remains constant for the first four days of cultivation. After five days the portion of non-fluorescent cells increases throughout the analysed samples (Tab. S1).

**Fig. 4:**
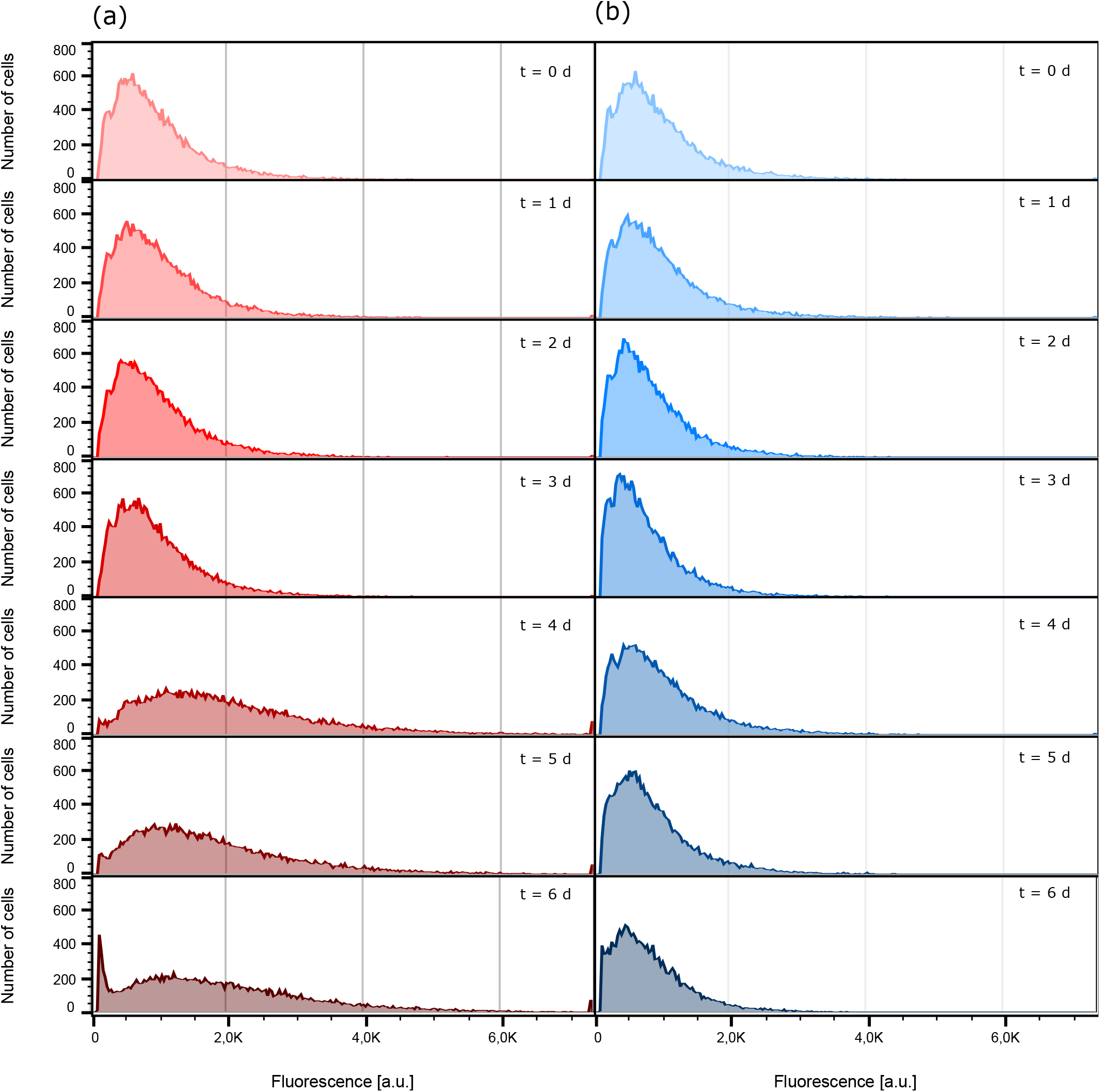
Comparison of eGFP fluorescence development. For illustrative reasons cells identified as non-fluorescent are not displayed. (a) Fluorescence intensity distribution inside the bioreactor at different sampling times. (b) Fluorescence intensity distribution inside the shake flask at different sampling times.

The data presented in Figure 4 only shows the population’s status at one specific timepoint during the cultivation. Therefore, these measurements are limited to population dynamics between distinct sampling times and do not yield any insights into dynamic fluorescence development of single cells. Furthermore, it is not possible to retain the same group of individual cells across the course of the cultivation using flow cytometric analysis, meaning that at every sampling time different cells are analysed. Yet, knowing if single cells show steadily increasing fluorescence levels, representing a constant product formation, or fluctuate in their productivity is a valuable information to classify the performance of a bioprocess. For this purpose, MSCC needs to be applied to examine single-cell dynamics.

Additional to performed population analysis, which shows highly comparable characteristics in general fluorescence distribution to the data obtained from bioreactor and shake flask via flow cytometry (Fig. S6), we investigated single-cell fluorescence development for a representative isogenic population to exemplify feasible single-cell analysis (Fig. 5a). Considering the doubling time of single cells, already after the first cell division the two originated daughter cells greatly differ in the duration until their second cell division by the factor of two (Fig. 5b). Looking at the respective video (Video S2) it appears to be very likely that this divergence arises from the asymmetrical division of the mother cell. In the following subsequent cell divisions both daughter cells’ offspring’s show only little variations (Fig. 5b).

**Fig. 5:**
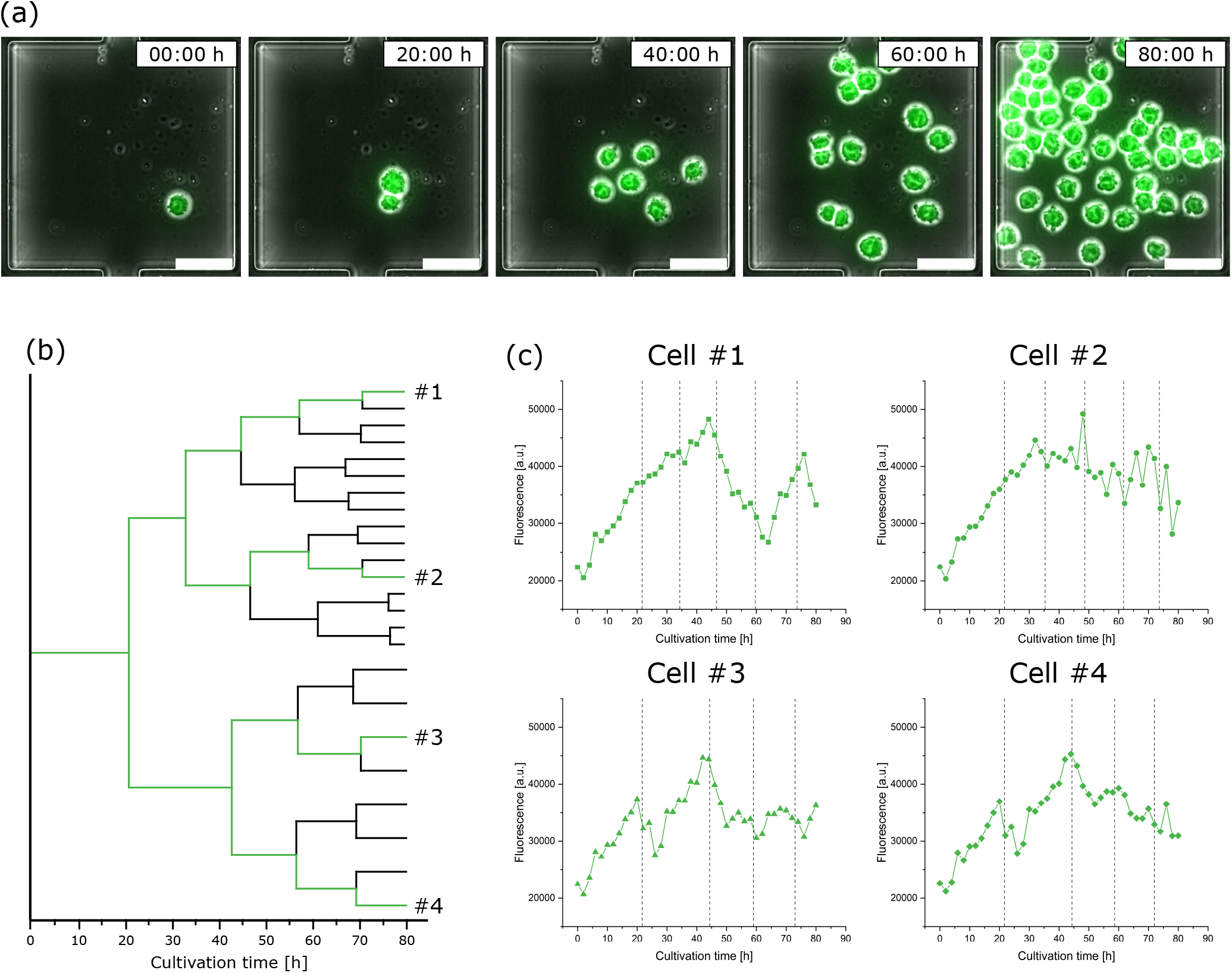
Analysis of eGFP-fluorescence single-cell dynamics during microfluidic cultivation. (a) Time-lapse image sequence showing the growth and fluorescence development of an isogenic microcolony (scale bar = 50 µm). (b) Lineage tree of the same isogenic microcolony. (c) Fluorescence development of four exemplary single cells over the cultivation time; dotted lines indicate cell division events.

This heterogenous behaviour cannot only be found in relation to growth but also has a noticeable impact on single-cell fluorescence. Figure 5c shows the fluorescence development of four exemplary cells over time, each from a different branch of the population’s lineage tree, with their respective cell division events marked by dotted lines. Looking at cell #1, varying stages of fluorescence increase and decrease can be identified. Although having the closest relation, the fluorescence course of cell #2 shows a slight but steady decrease and does not feature comparable fluctuating tendencies like cell #1. Comparing cell #3 against cell #4 both vary only slightly in their course after originating from a common progenitor cell.

Relating the sections of fluorescence increase and decrease from Figure 5c to the respective moments of cell division no obvious interrelationship can be identified. Nevertheless, the already discussed asymmetrical division of the initial not only influences following cell division events but also results in a drastic fluorescence decrease of one daughter cell.

Disregarding the differences in their individual fluorescence development, all cells in Fig. 5c do not show a steady increase of fluorescence intensity over the whole cultivation course like the population analysis via flow cytometry in case of bioreactor and shake flask cultivation might indicate (Fig. 4). This observation stresses the importance of dynamic single-cell analysis since detecting fluctuating production behaviour represents the first step towards eventually engineering bioprocesses for higher productivities by more stable product formation behaviour.

## Conclusion and outlook

Environmental control, live cell imaging, and high spatio-temporal resolution make MSCC a highly valuable miniaturisation tool, as single-cell dynamics under constant cultivation conditions can be analysed over multiple generations and therefore intercellular differences in growth behaviour or fluorescence-coupled protein expression can be investigated. These analyses are not performable applying standard average measurements as it is common with other small-scale systems like microtiter plates or miniaturised bioreactors.

The study presented here shows that MSCC-generated data is comparable to data from lab-scale cultivation approaches in all investigated aspects namely growth, cellular morphology, and production behaviour. Regarding growth behaviour, cells cultivated on-chip showed the same growth rate as populations cultivated in shake flasks or bioreactors. Likewise, cellular morphology concerning single-cell diameter of MSCC was representative for cells that were cultivated in bigger scales. Despite the different quantification approaches, for eGFP production the fluorescence distributions throughout the analysed populations were comparable as well. Additional to the population dynamics investigated via flow cytometry, MSCC allowed analysis of single-cell fluorescence dynamics and revealed phases of fluorescence increase and decrease over the course of cultivation which would have stayed hidden applying conventional flow cytometric analysis.

The proven comparability between the microfluidic miniaturisation tool and shake flask or lab-scale bioreactor cultivation enables MSCC to be applied for numerous applications in basic research as well as for bioprocess development. In the context of mammalian bioproduction, especially studying cellular heterogeneity concerning growth behaviour and productivity inside isogenic populations is of utmost interest since it can have a severe influence on bioprocess robustness and outcome (Paul and Herwig 2020). Particularly the number of generations and the resulting cellular heterogeneity, with its effects from single-cell cloning up to commercially application, represents a very important question in process development (Frye et al. 2016; Rugbjerg and Sommer 2019), that eventually can be analysed more closely by the means of MSCC.

## Acknowledgements

The authors gratefully acknowledge support by the “European Regional Development Fund (EFRE)” through project “Cluster Industrial Biotechnology (CLIB) Kompetenzzentrum Biotechnologie (CKB)” (34.EFRE-0300095/1703FI04) for Oliver Hertel. Furthermore, we thank Melinda Go for kindly providing the eGFP-producing CHO-K1 cell pool.

## Notes

### Competing Interest Statement

The authors have declared no competing interest.

