## Supplementary information for "Growth and eGFP-production of CHO-K1 suspension cells cultivated from single-cell to lab-scale"

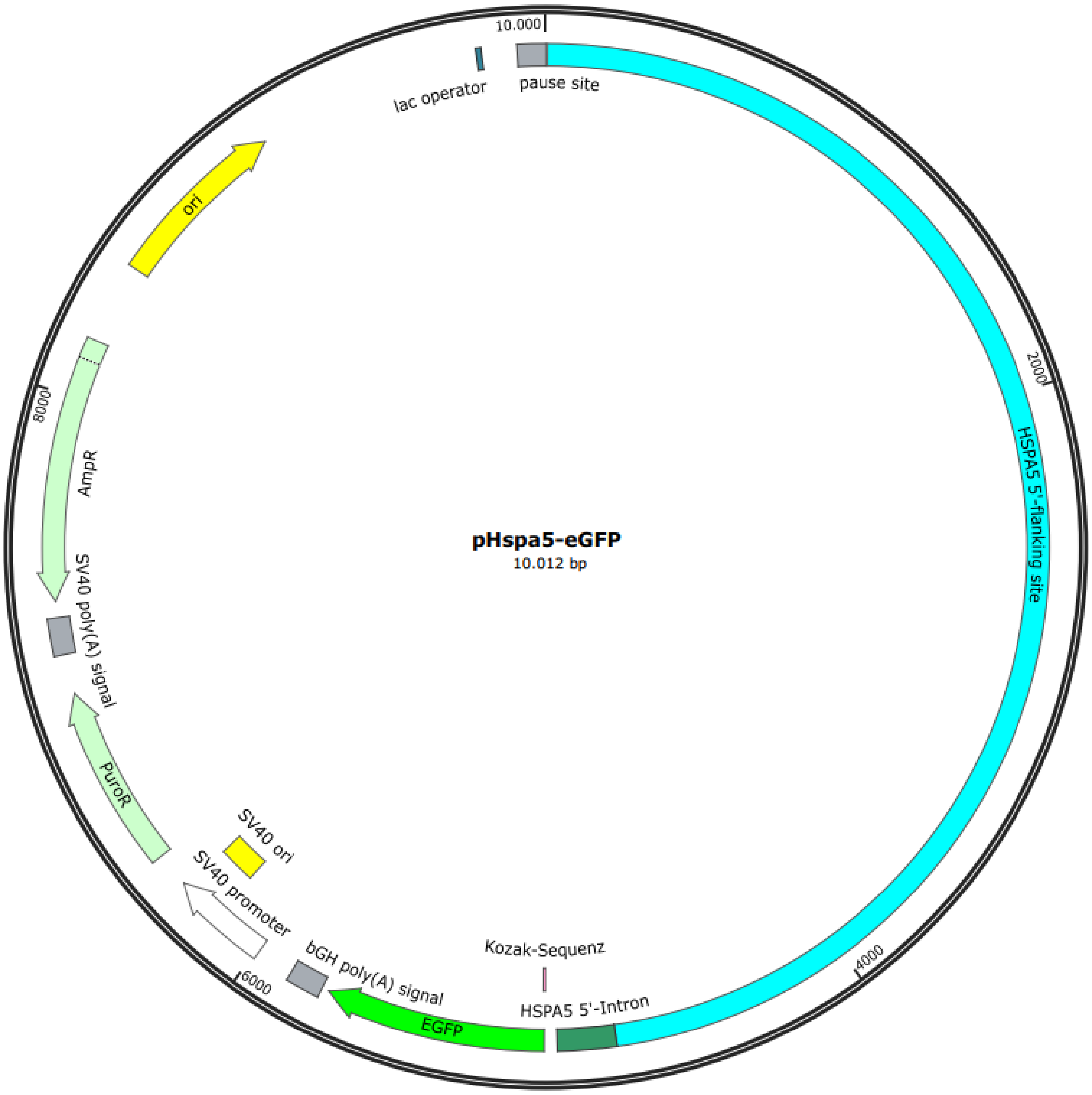

Fig. S1: Transfection vector showing the endogenous HSPA5 promoter and eGFP gene as well as a puromycin resistance for selection. The map was created using SnapGene® (GSL Biotech LLC, USA).

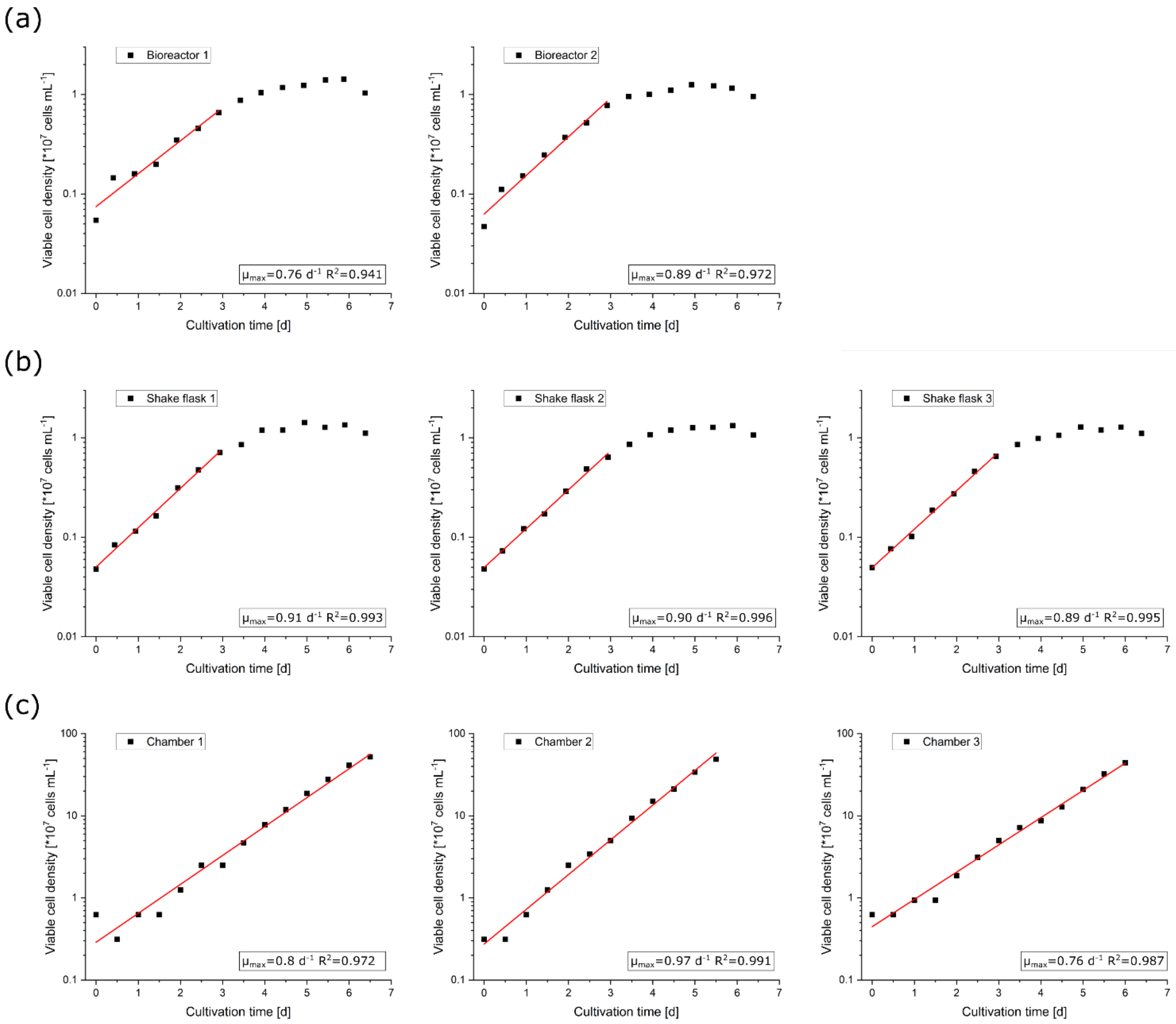

Fig. S2: Semi-logarithmically plotted growth profile of the (a) bioreactors, (b) shake flasks, and (c) MSCCs with their resulting linear fit in exponential growth phase. For every colony the graphically determined growth rate μ with its coefficient of determination $R^2$ are placed in the respective plot at the bottom right.

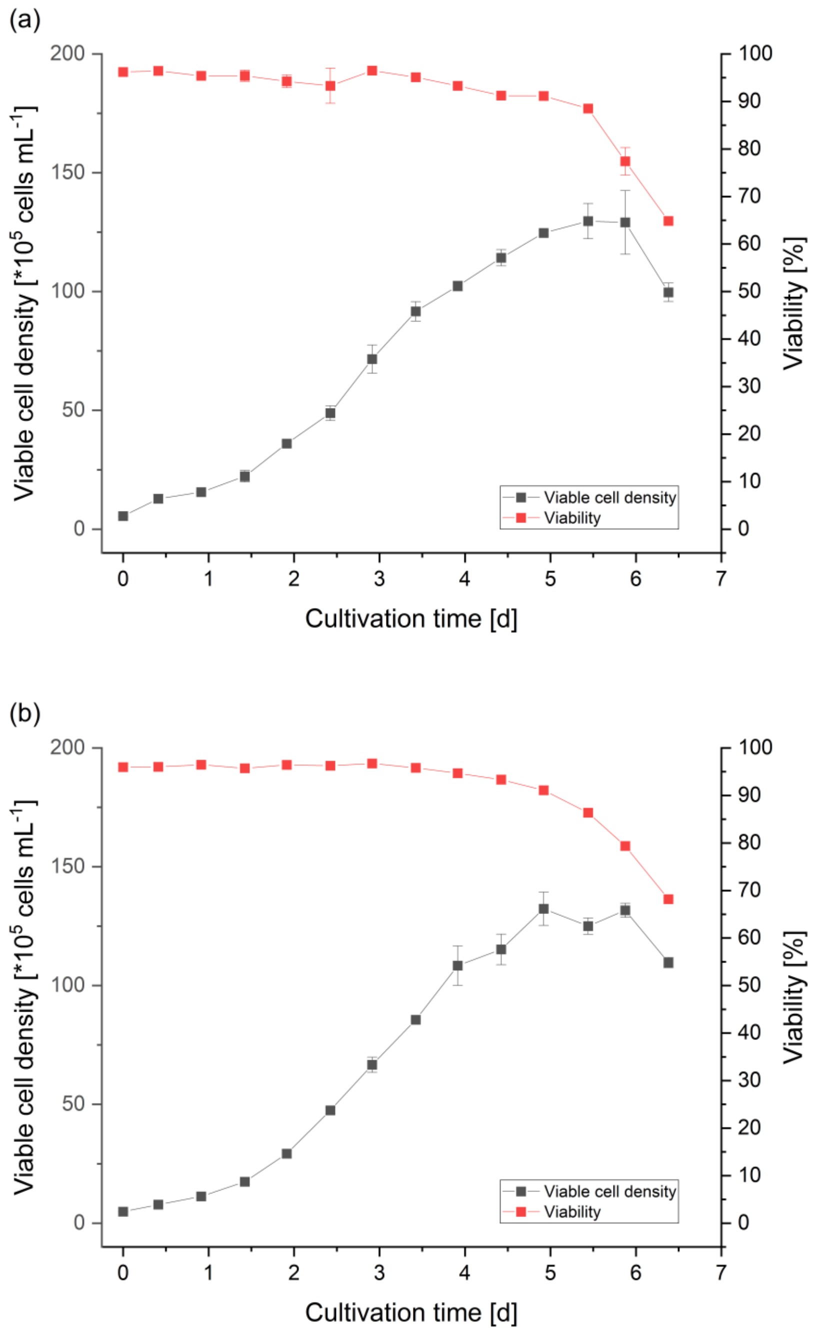

Fig. S3: Viable cell density and viability profile of the (a) bioreactor and (b) shake flask cultivation. Both viable cell density and viability are averaged for the respective replicates (bioreactor $n_{cultivation}$ = 2, shake flask $n_{cultivation}$ = 3).

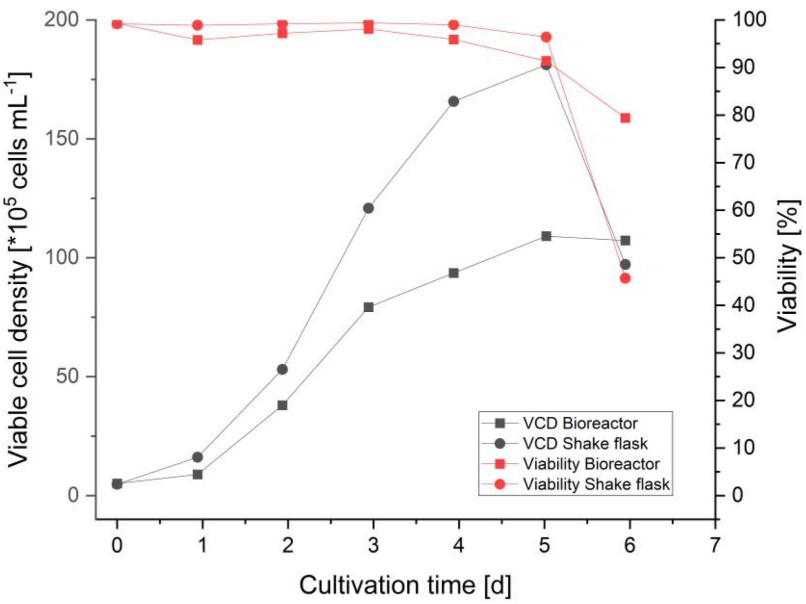

Fig. S4: Viable cell density (VCD) for bioreactor and shake flask cultivation of the eGFP-producing CHO-K1 cell pool.

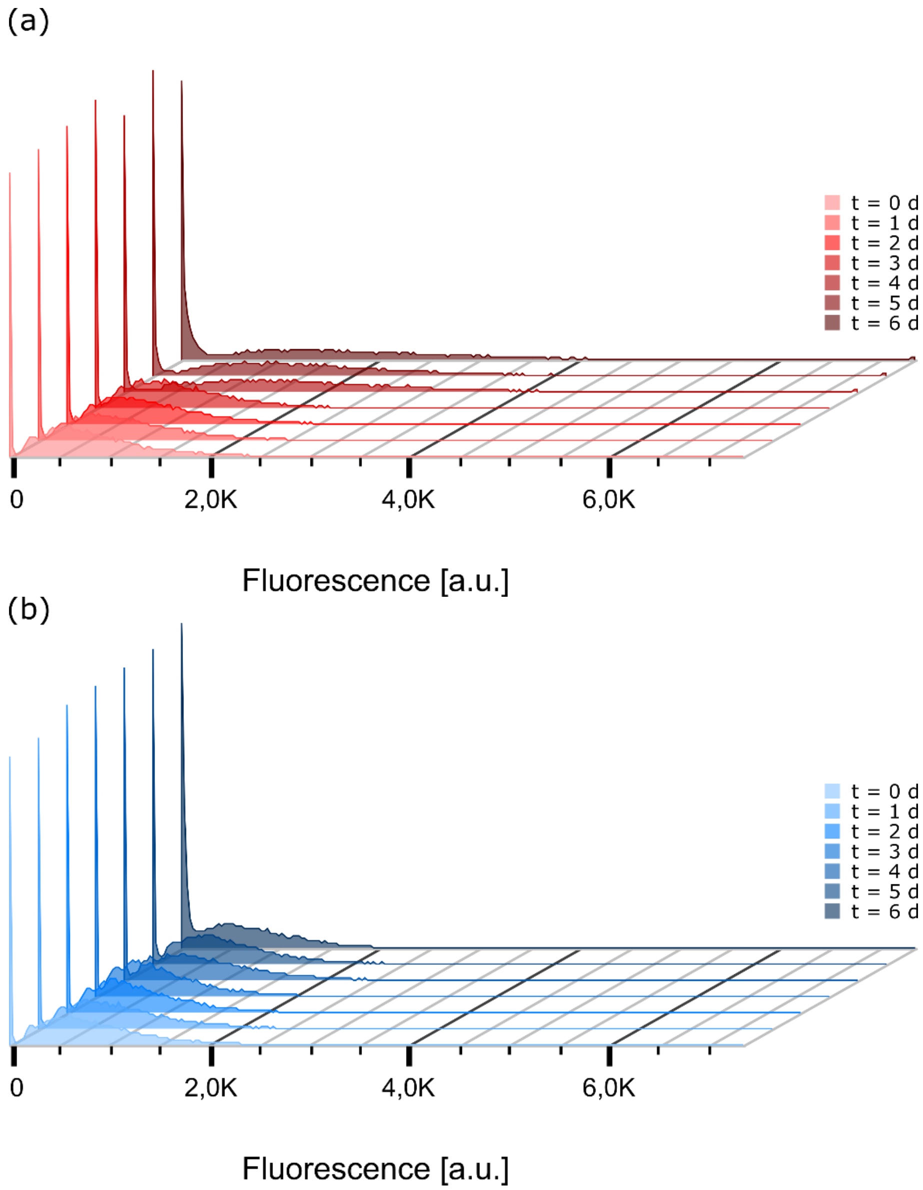

Fig. S5: eGFP distribution at different sampling times during (a) bioreactor and (b) shake flask cultivation. Displayed are both fluorescent and non-fluorescent cells of the respective sample.

Tab. S1: Summary of the flow cytometric analysis of the eGFP-production from shake flask and bioreactor illustrated in Fig. S6. For every sampling time the number of analysed cells, the percentage of non-fluorescent cells, the percentage of fluorescent cells, and the median of the fluorescence intensities are displayed.

| Sampling time | Parameter | Shake flask | | Bioreactor | |
|---|---|---|---|---|---|
| | | Percentage / % | Cells / - | Percentage / % | Cells / - |
| t0 | Singularised | 99.4 | 29830 | 99.6 | 29874 |
| | eGFP negative | 20.8 | 6194 | 20.4 | 6099 |
| | eGFP positive | 79.2 | 23633 | 79.6 | 23770 |
| | Median of fluorescence [a.u.] | | 804 | | 799 |
| t1 | Singularised | 99.7 | 29899 | 99.5 | 29864 |
| | eGFP negative | 21 | 6272 | 21.4 | 6386 |
| | eGFP positive | 79 | 23627 | 78.6 | 23479 |
| | Median of fluorescence [a.u.] | | 799 | | 858 |
| t2 | Singularised | 99.8 | 29937 | 99.7 | 29915 |
| | eGFP negative | 22.3 | 6674 | 22 | 6578 |
| | eGFP positive | 77.7 | 23265 | 78 | 23338 |
| | Median of fluorescence [a.u.] | | 704 | | 819 |
| t3 | Singularised | 99.8 | 29947 | 99.9 | 29957 |
| | eGFP negative | 22.9 | 6848 | 22.4 | 6725 |
| | eGFP positive | 77.1 | 23100 | 77.6 | 23234 |
| | Median of fluorescence [a.u.] | | 638 | | 817 |
| t4 | Singularised | 99.8 | 29951 | 97.7 | 29314 |
| | eGFP negative | 22.9 | 6863 | 21.5 | 6314 |
| | eGFP positive | 77.1 | 23088 | 78.5 | 23000 |
| | Median of fluorescence [a.u.] | | 822 | | 1744 |
| t5 | Singularised | 99.9 | 29974 | 99.7 | 29910 |
| | eGFP negative | 24.1 | 7213 | 24.6 | 7345 |
| | eGFP positive | 75.9 | 22762 | 75.4 | 22566 |
| | Median of fluorescence [a.u.] | | 727 | | 1528 |
| t6 | Singularised | 99.9 | 29957 | 99.7 | 29917 |
| | eGFP negative | 37.8 | 11309 | 29.3 | 8768 |
| | eGFP positive | 62.3 | 18649 | 70.7 | 21153 |
| | Median of fluorescence [a.u.] | | 706 | | 1664 |

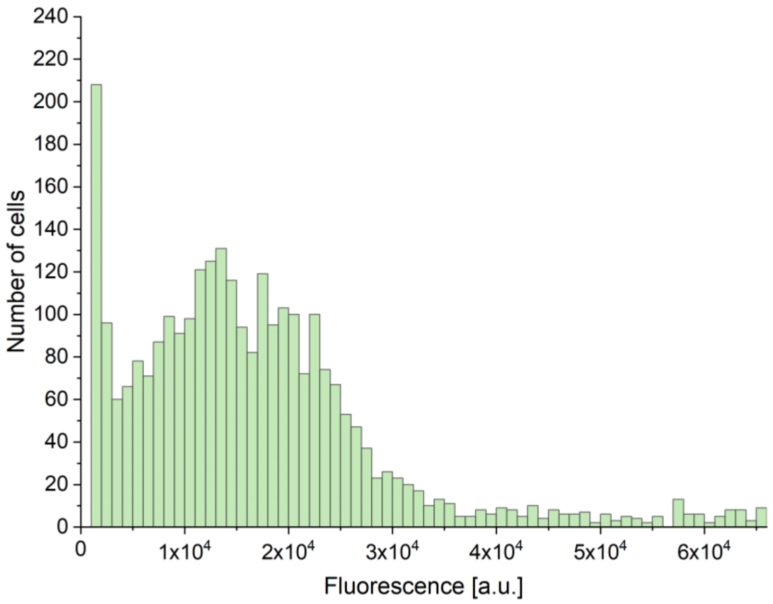

Fig. S6: eGFP distribution of 44 analysed microcolonies after 80 h (during exponential growth) of microfluidic single-cell cultivation ($n_{cells}$ = 2,800).
